# Functional Role of Annexins in Zebrafish Caudal Fin Regeneration – A Gene Knockdown Approach in Regenerating Tissue

**DOI:** 10.1101/342238

**Authors:** Mir Quoseena, Sowmya Vuppaladadium, Shahid Hussain, Swarna Bharathi, Mohammed M Idris

## Abstract

Regeneration is an adaptive phenomenon with wide biological implications spread heterogeneously in almost all the organism including human beings. The ability of regeneration varies from species to species for its impediment. Epimorphic regeneration of zebrafish caudal fin tissue is most widely studied regeneration mechanism for its discrete and rapid regenerative capability. Several genes/proteins were found to be associated with zebrafish caudal fin tissue regeneration. Here, we have evaluated the functional role of *Annexin 2a* and *2b* genes in adult zebrafish caudal fin tissue undergoing regeneration using a novel CRSISPR-Cas9 gene knock down approach. Knock down of both the genes individually elicited in decelerated regeneration and down regulation of target genes and its partner genes/proteins such as ANXA1a, ANXA5b and ANXA13. This study validates a novel gene targeting approach and the possible role of annexin in regeneration mechanism.

---

The eukaryotic annexin protein family is the largest calcium-binding group that lacks E-F hand^1^. It consists of eleven proteins that have been identified in vertebrates (ANXA1–7, 9, 11, 13, and 31). Four genes (anx1a, anx2a, anx5, and anx11a) were identified in zebrafish^2^. All ANXAs show a highly conserved intron–exon organization^1^ and share a characteristic, highly conserved 70-amino acid domain that is repeated 4 to 8 times, with an ability of calcium dependent anionic phospholipids binding (with the exception of ANX31, which lacks the calcium binding site^3^). The function of this evolutionarily conserved family of proteins remains poorly understood, though ANXA5 is being widely studied for live apoptotic imaging in zebrafish^4^. Evolutionary history of this gene family was examined through many studies^5^. ANX2a and 2b are highly expressed in zebrafish regeneration tissue^6^.

1970 marked the beginning of new era for biology, recombinant DNA technology gave molecular scientist the ability to manipulate DNA. CRISPR (clustered regularly interspaced short palindromic repeats) - Cas (CRISPR-associated genes) RNA-guided nucleases, an adaptive immunity system found in bacteria and archaea is one of the simplest platform of gene manipulation identified^7^ till date. According to Chylinski *et.,al.* CRISPR-Cas marks fragments of DNA from its previously encountered viruses and plasmids as an array of CRISPR repeats and are later used as guide CRIPSR RNA (crRNA) to target the kindred new infections^8, 9^.

The Type II CRISPR system of Streptococcus pyogenes is used by various bacteria as a mechanism to inactivate foreign DNAs^10^. Several reports have also established knock in zebrafish lines with this technology^11-14^. Its ease of use, low cost, and the possibility of multiplexing make it a prime tool for large-scale reverse genetic screens in zebrafish^15^. In many cases, the investigation of gene *function in vivo* requires the spatiotemporal control of gene silencing. Here, we present the rationale for tissue specific gene inactivation in zebrafish using the CRISPR/Cas9 technology and detailed methodology to achieve it.

On the basis of our earlier work we have identified annexin genes responsible for regeneration in zebrafish using transcriptomics. We decided to do further study on Annexin gene families as there is more than 70% homology between zebrafish and human annexins. To study the effect of these target genes we had designed current study to Knock down ANXA2a and 2b genes using CRISPR and study its effect at phenotypic as well as genotypic levels.

## Results

### Synthesis of target and Cas9 mRNA

A total of 7 ANXA2a and 2 ANXA2b specific targets of 22bp length having BSaI restriction site were designed using the ZiFiT program (Table 1, Figure 1a). All the targets were designed to be in the exonic region of the gene. The targets were synthesized as oligonucleotides and dimerized with their respective antisense oligonucleotide and cloned in the pDR274 vector (Figure 1b). RNA specific to the target site were synthesized from the positive and confirmed clones. The mRNA obtained from each target site were pooled for respective target genes. Similarly, Cas9 mRNA were synthesized from the pMLM3613 vector (Figure 1b) and mixed with pooled ANXA2a and 2b specific target mRNA and electroporated to the animals after amputation.

**Table 1:**
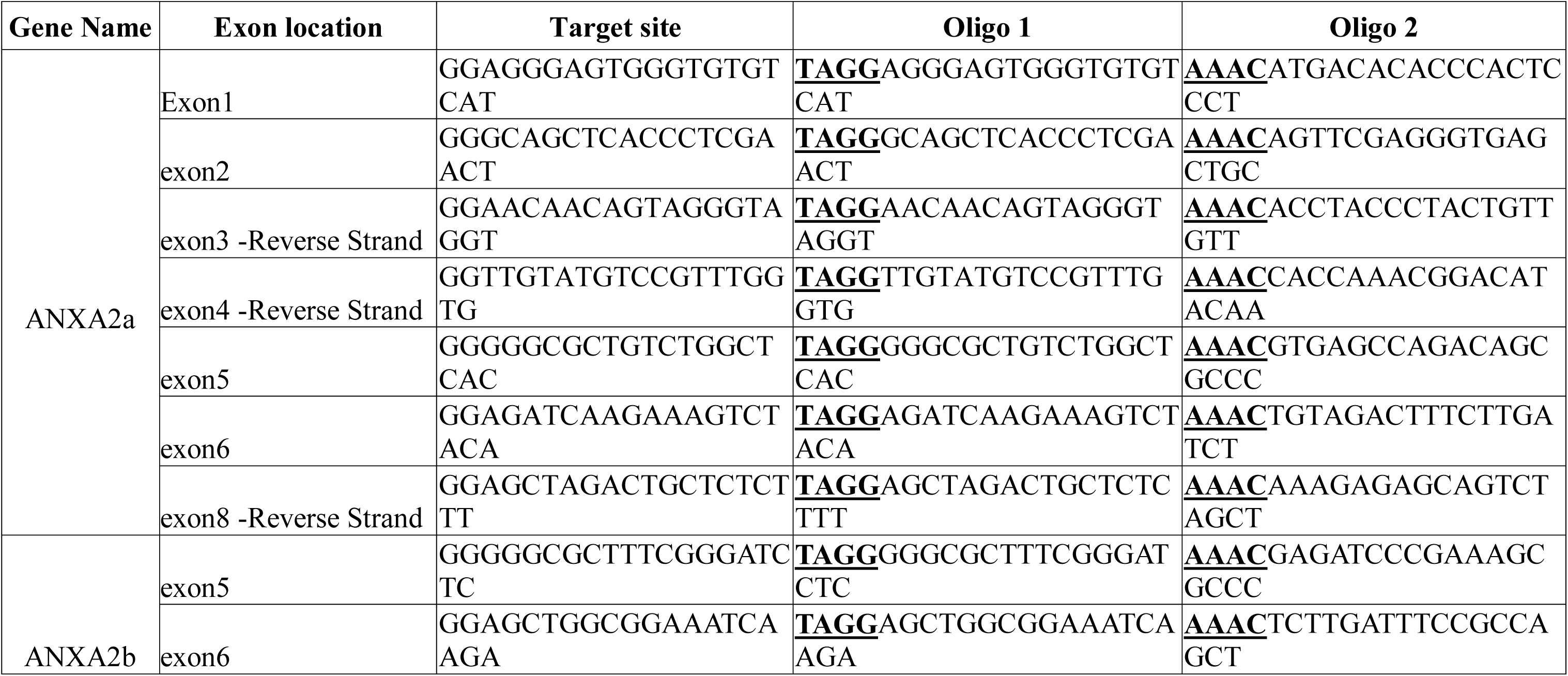
List of CRISPR targets specific to ANXA2a and 2b genes and its forward and reverse oligonucleotide sequences.

**Figure 1:**
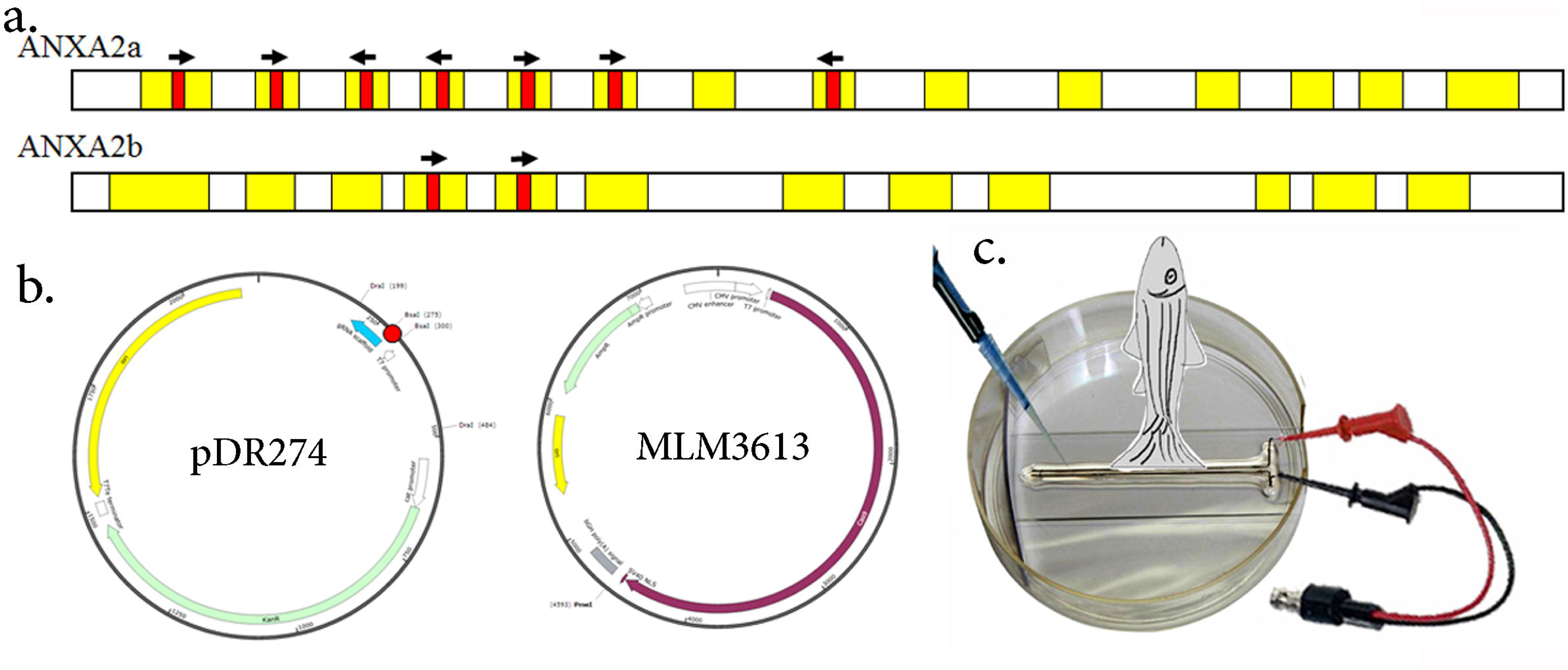
1a. Schematic map of ANXA2a and 2b gene with exon (shaded in yellow) and CRISPR-Cas Target site (shaded in red) with arrow indicating the orientation of the CRISPR-Cas target. b. plasmid map of DR274 with red dot indicating the cloning site and MLM3613 vectors. c. Cartoon detailing the electroporation protocol of CRISPR-Cas9 RNA in to the adult zebrafish amputated fin tissue using electroporator.

### Phenotype analysis

Based on phenotype analysis of the zebrafish caudal fin tissue post amputation and electroporation it was found that there is a blatant effect in the growth of the fin tissue due to CRISPR target (Figure 2). The structural regeneration of the fin tissue was found significantly reduced with retarded regeneration growth in both ANXA2a and 2b targeted fin tissue from 5dpa onwards (Figure 2b). Comparison of the growth of fin length showed a strong association of retarded growth in Annexin gene targeted fin tissue against the control and CAS9 targeted regenerating tissues. Also, the shape of the regenerating fin tissue was lost due to ANXA2a and 2b targets in the later time points post amputation (Figure 2a).

**Figure 2:**
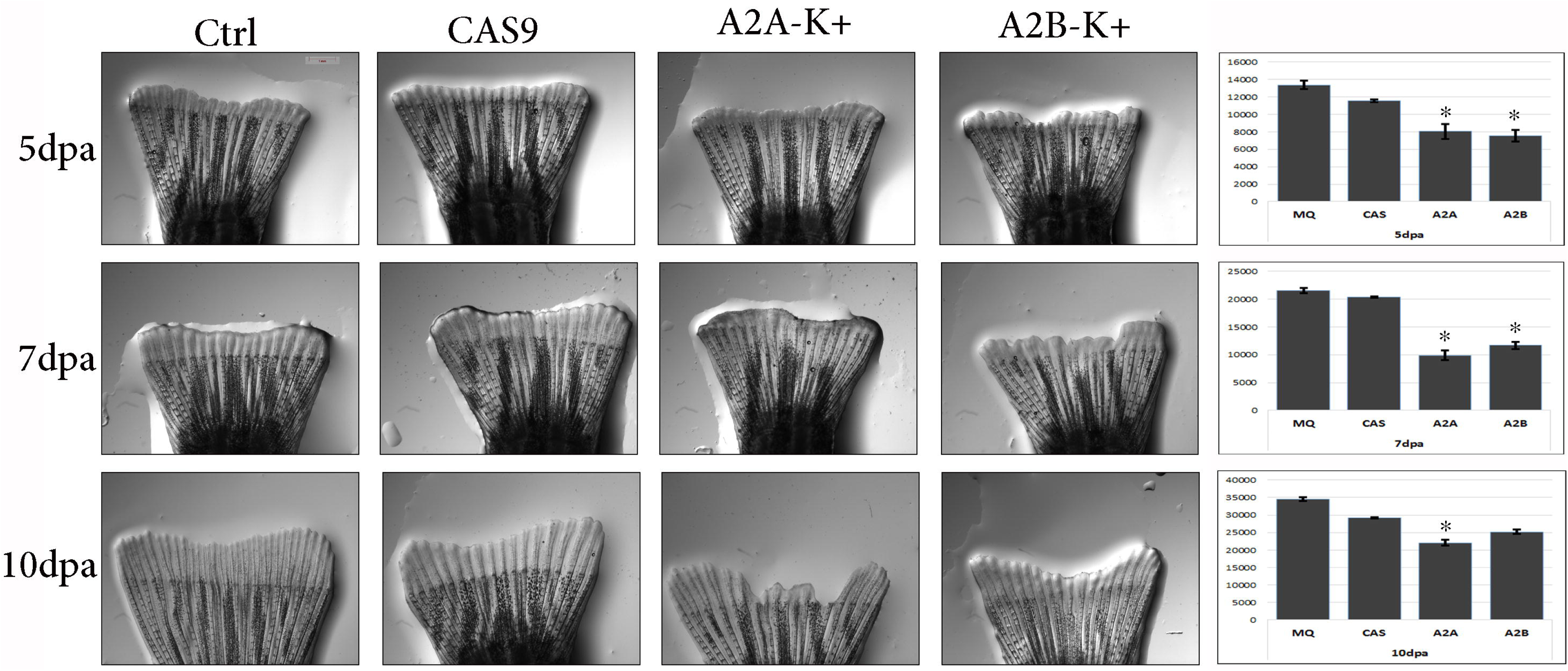
2a. Phenotype of regenerating caudal fin tissue at 5, 7 and 10 dpa electroporated with MQ, Cas9, ANXA2a and ANXA2b CRISPR targets. B. Bar diagram detailing the rate of fin tissue growth at respective time points and electroporation. The fin growth was represented as mean +SEM; * p<0.05 vs MQ-Ctrl group. (Student t-test, N=3).

### Genotype analysis

Expression of ANXA2a and 2b genes were found to be down regulated in the respective gene knock down fin tissue against CAS9 and MQ control tissues at 1, 2 and 7 dpa (Figure 3a). The expression of ANXA2a gene amplified by target region specific primer showed a complete reduction in the gene expression or a truncated PCR amplicon, validating the positive CRISPR gene action (Figure 3a, 3b). Similarly, ANXA2b gene amplification was also found down regulated in the ANXA2b targeted fin tissues. The expression of genes was found to be recovered at 7dpa in both the gene targeted tissues. ANXA2a showed more effect than ANXA2b gene in comparison to control and normal fin tissues gene expression. Interestingly the expression of ANXA2b was also found down regulated in the ANXA2a targeted fin tissue and vice versa. Expression analysis of other annexin family genes such as ANXA1a, 5b and 13 also found to be down regulated in the ANXA2a and 2b knock down tissues at 1 and 2 dpa. A full recovery of the gene expressions was found at 7dpa for all the annexin genes (Figure 3a, 3b). The expression of ANXA2b did not recover as like ANXA2a in ANXA2b KD+ tissue at 7dpa. The expression of ANXA2a and 2b were found either down regulated or expressed as truncated amplicon upon knockdown of the ANXA2a and 2b genes confirming the action of the CRISPR on target.

**Figure 3:**
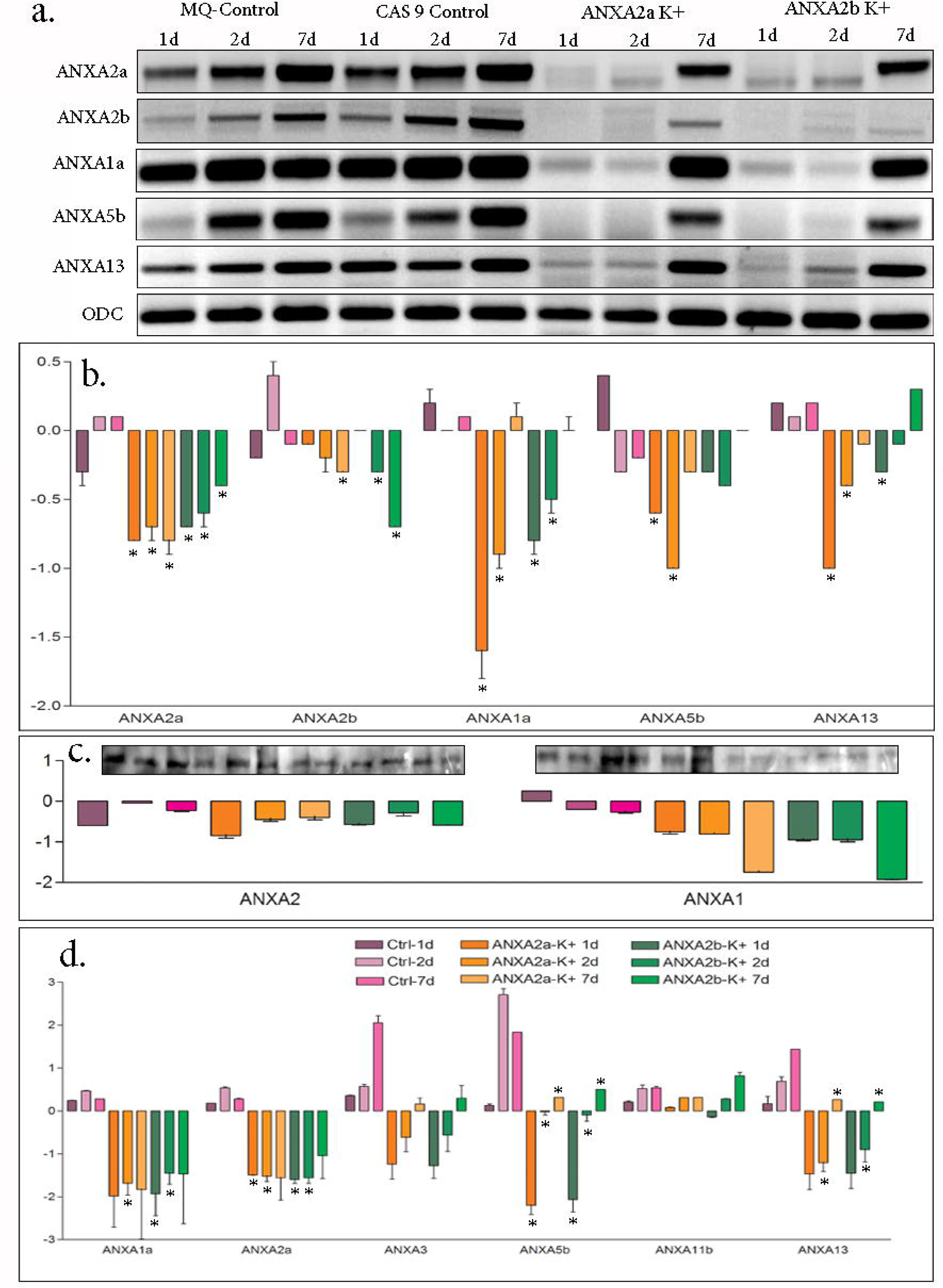
a. Annexin gene expression based on PCR and gel electrophoresis for MQ-Ctrl, CAS9 Ctrl, ANXA2aK+ and ANXA2bK+ fin tissue samples; b. Band intensity and bar graph of expression of ANXA2a, 2b, 1a, 5b and 13 genes. c. Western blot image of ANXA2 and ANXA1 proteins and its expression intensity. d. Expression of Annexin proteins based on iTRAQ based Quantitative proteomic analysis. The expression of genes and proteins were represented as mean +SEM; * p<0.05 vs MQ-Ctrl group. (Student t-test, N=3).

### Proteomic Analysis

Proteomic analysis involving western blot assay and quantitative proteomic analysis of the regenerating caudal fin tissue targeted with ANXA2a and 2b CRISPR showed a significant down regulation of the ANXA2 and ANXA1 proteins along with other Annexin family proteins. Western blot analysis for ANXA1 and ANXA2 protein (both ANXA2a and 2b) expression in the regenerating fin tissue targeted with ANXA2a and 2b CRISPR showed a non-significant down regulation of both the proteins (Figure 3c). iTRAQ based Quantitative proteomic analysis of protein fractions matching to annexin family proteins showed significant down regulation of ANXA1a, ANXA2a, ANXA5b and ANXA13 proteins (Table 2, Figure 3d) at 1, 2 and 7 dpa. ANXA11b expression was not found to be modified due to knock down of ANXA2a and 2b genes as like other annexins (Figure 3d).

**Table 2:** Expression of Annexin family proteins in control, ANXA2a and ANXA2b targeted zebrafish caudal fin tissues based on iTRAQ based quantitative proteomics. The expressions were shown as mean of replicates compared against MQ-Ctrl samples of respective time points.

| Gene Name | Ctrl-1d | Ctrl-2d | Ctrl-7d | ANXA2a-K+<br>1d | ANXA2a-K+<br>2d | ANXA2a-K+<br>7d | ANXA2b-K+<br>1d | ANXA2b-K+<br>2d | ANXA2b-K+<br>7d |
| --- | --- | --- | --- | --- | --- | --- | --- | --- | --- |
| ANXA1a | 0.24 | 0.47 | 0.28 | -1.98 | -1.69 | -1.83 | -1.93 | -1.45 | -1.47 |
| ANXA2a | 0.18 | 0.54 | 0.28 | -1.49 | -1.52 | -1.56 | -1.60 | -1.56 | -1.04 |
| ANXA3 | 0.36 | 0.57 | 2.05 | -1.24 | -0.61 | 0.16 | -1.28 | -0.56 | 0.30 |
| ANXA5b | 0.13 | 2.71 | 1.84 | -2.20 | -0.02 | 0.32 | -2.07 | -0.09 | 0.50 |
| ANXA11b | 0.21 | 0.52 | 0.54 | 0.08 | 0.31 | 0.32 | -0.14 | 0.28 | 0.82 |
| ANXA13 | 0.17 | 0.69 | 1.44 | -1.47 | -1.21 | 0.26 | -1.45 | -0.90 | 0.21 |

## Discussions

Here, a novel technique based on *in vivo* tissue specific electroporation was developed and optimized to improve the efficiencies of genome editing of genes by CRISPR in zebrafish by using a CRISPR/Cas9 system. Annexin genes are 70% conserved and ANXA2b is the only member of family absent in humans. We have shown direct association of ANXA2a and 2b with fin regeneration in zebrafish^6, 16.^ The low efficiencies of tissue specific genome editing are the common issues confronted with many researchers. To solve this problem, different approaches were developed and evaluated. In this study, the tissue specific genome editing with no mismatch targeting for all the replicates was confirmed, which indicated that the electroporation of Cas9 capped RNAs into tissue could reduce the time for Cas9 translation, and subsequently improve the probability of mutations transferred to the next generation. This technique of electroporation following amputation has made gene targeting much easier than the complicated microinjection-based protocol.

Thus, this novel technique based on *in vivo* tissue specific electroporation can improve the efficiencies of knock-out, knock-in and germline transmission by using a CRISPR/Cas9 system. It’s worth mentioning that an increasing efficiency of knocking out ANXA2a and 2b was achieved in this study. This has an effect on the expression levels of other members of the gene family, whether the knocking out efficiency was correlated between different genes of annexin family deserves further studies. The choices of effective target site and appropriate time points were the critical factors to affect the probability of the successful gene editing.

In conclusion, a new approach based on *in vivo* tissue specific gene electroporation was developed and evaluated in this study. This technique was confirmed to improve the efficiencies of genome editing in zebrafish by using a CRISPR/Cas9 system. This novel strategy will reduce the unnecessary labor-intensive screening and will significantly increase the speed of generating desirable mutants in zebrafish. This study also confirms that ANXA2a and ANXA2b are the important genes involved in regeneration and it has a correlated role along with ANXA1, ANXA5 and ANXA13 gene for the regeneration of caudal fin tissue in zebrafish.

## Methods

### Construction of Expression Vector with sgRNA

CRISPR sgRNA targets specific to ANXA2a and ANXA2b genes were designed using ZiFiT targeter program. All the targets were synthesized as oligonucleotides and anti-sense oligonucleotides (Bioserve, India) (Figure 1a). Oligonucleotides pairs were annealed to each other by incubating 25 pmoles of forward and reverse oligonucleotides at 95°C for 3 mins followed by touch down cooling of the mix to 30°C. pDR274 vector obtained from Addgene (plasmid #42250 ^17^), was used as the expression vector for the synthesis of sgRNA specific to the ANXA2a and ANXA2b targets. 100 ng pDR274 vector was linearized with BsaI and ligated with annealed oligos using RAPID ligase (Invitrogen), transformed in DHα cells using heat shock transformation protocol and selected the positive clone in LB kanamycin selection plate (Figure 1b). Colony PCR and sequencing of the positive clones were performed using M13F and gene specific primers.

### sgRNA mRNA synthesis

sgRNA mRNA specific to ANXA2a and ANXA2b targets were synthesized following DraI restriction digestion for eluting 282 bp fragment containing the T7 promoter and the sgRNA targets. sgRNA has been transcribed by *in vitro* transcription using the MAXI script ® T7 kit (Thermo Fischer Scientific) as per manufacturer’s protocols. DNase treated sgRNA was then purified and precipitated by acidic phenol chloroform and alcohol precipitation. The dissolved RNA was aliquoted into multiple tubes and then was quantified using a spectrometer. Integrity of sgRNA was confirmed by gel electrophoresis.

### Cas9 mRNA synthesis

Cas9 mRNA was synthesized from pMLM3613 vector obtained from Addgene (plasmid #42251^17^), through linearizing the vector with PmeI restriction enzyme, gel purification and reverse transcriptase synthesis using mMESSAGE mMACHINE ^®^ 7 Ultra kit (Thermo Fischer Scientific) (Figure 1b). The synthesized Cas9 mRNA were treated with DNaseI to remove vector DNA followed by polyA tailing using E-PAP enzyme. Poly A tailed cas9 mRNA were precipitated using Lithium Chloride and dissolved in nuclease-free water.

### Amputation and Electroporation

9 to 12-month old Adult zebrafish maintained under standard laboratory conditions were selected, anesthetized and distal halves of the caudal fins were amputated under sterile condition^16^. Cas9 and sgRNA mRNA were electroporated to the amputated fin tissue using a novel approach. 1:5 ratios of sgRNA (50 ng/µl) and Cas9 mRNA (250 ng/µl) were mixed prior to electroporation and 25 µl of the mix were placed between the two round wire electrodes (BTX) in the glass micro slides. Amputated zebrafish caudal fin tissue was placed in the space between the electrodes vertically by holding the anaesthetized fish in a sponge such that the tip of the fin touches the RNA solution during electroporation. Electroporation was performed on the fin tissue having a set 50V for 30ms pulse length and 3 pulse repeats in the electroporator (ECM2001, BTX Harvard apparatus, USA) (Figure 1c). Electroporation was performed immediately after amputation, as first dose and 24 hrs post amputation as second dose. For 1dpa genotype analysis only one dose of electroporation was given. Control experiments were performed for each set of experiment by electroporating only Cas9 mRNA and milli Q water. All the experiments were performed in replicates (n=3). The procedure of animal experiments and the gene manipulation has been performed in accordance with the protocol approved by the Institutional animal ethics committee of Center for Cellular and Molecular Biology (IAEC/CCMB/Protocol # 50&52/2013).

### Genotype and Phenotype analysis

For phenotype analysis, fin tissues were collected at 5, 7 and 10-day post amputation and electroporation. The regenerated fin tissue growth was measured using image J software. For genotype analysis the tissues were collected at 1, 2 and 7-day post amputation and electroporation. RNA was extracted from the individual fin tissues using TRI reagent (Sigma, USA)^16^. cDNA was synthesized from 100 µgms of total RNA samples extracted from individual fin tissues using Reverse Transcriptase Core Kit (Eurogentec, Belgium). Validation and expression of the targeted gene, ANXA2a and 2b genes and non-targeted genes ANXA1a, ANXA5b, ANXA13 and ODC were measured based on quantitative RTPCR on the collected individual fin tissue RNA using gene specific primers^6^. The expression of the genes was compared against control time point samples.

### Proteomic and WB analysis

iTRAQ based quantitative proteomic analysis was performed on the fin tissues collected after 1, 2 and 7-day post amputation and electroporation. The protein was extracted from the pooled tissue of each stage, quantified and 1DE gel electrophoresed. The gels were fractioned for different molecular weight ranges and In-gel trypsin digestion was performed. Cas9 control, ANXA2a and ANXA2b gene of targeted time point samples were labelled with iTRAQ labels 114, 115, 116 and 117 respectively^18^. MS/MSMS analysis of the peptide samples corresponding to the ANXA2a and 2b molecular weight fraction were analyzed in Q-Exactive Mass spectrometer. The expression of the protein was analyzed for its time point expression for 1, 2 and 7 dpa. Western blot analysis of the protein samples was also performed for ANXA2, ANXA1 and ODC protein expression using their respective mice specific antibody for the estimation of protein expression^6, 16^.

### Data Analysis

All the data obtained from various measurements were analyzed using MS-Excel and the statistical significance of the data was analyzed using PRISM6 software.

## Acknowledgments

The authors are thankful to Dr. Keith Joung (for gifting both the DR274 and MLM3613 vectors) and Dr. Gopal Kushwah for helping in getting the vector DNA. The work was supported by CSIR-YSA (Dr. Mohammed Idris) project fund. Authors are also thankful to Ms. Noorul Fowzia for critically reviewing the manuscript.

## Author contributions

MQ, SV, SH and SB performed the experiments and analyzed the data, MMI designed the experiment and wrote the manuscript.

## References

1. Smith, P. & Moss, S. Structural evolution of the annexin supergene family. Trends Genet. 10: 241–246 (1994).

2. Farber, S. D., De Rose, R. A., Olson, E. S. & Halpren, M. E. The zebrafish annexin gene family. Genome Res. 13: 1082–1096 (2003).

3. Moss, S. The annexins. In: The annexins (ed. S.E. Moss), pp. 1–10. Portland Press, Chapel Hill, NC (1992).

4. van Ham, T. J., Mapes, J., Kokel, D. & Peterson, R. T. Live imaging of apoptotic cells in zebrafish. FASEB J. 24: 4336–4342 (2010).

5. Morgan, R. O. & Pilar Frenandez, M. Distinct annexin subfamilies in plants and protists diverged prior to animal annexins and from a common ancestor. J. Mol. Evol. 44: 178–188 (1997).

6. Saxena, S. et al. Role of Annexin gene and its regulation during zebrafish caudal fin regeneration. Wound Repair Regen. 24: 551–559 (2016).

7. Deltcheva, E. et al. CRISPR RNA maturation by trans-encoded small RNA and host factor RNase III. Nature 471: 602–607 (2011).

8. Chylinski, K., Le Rhun, A. & Charpentier, E. The tracrRNA and Cas9 families of type II CRISPR-Cas immunity systems. RNA Biol. 10: 726–737 (2013).

9. Chylinski, K., Makarova, K. S., Charpentier, E. & Koonin, E. V. Classification and evolution of type II CRISPR-Cas systems. Nucleic Acids Res. 42: 6091–6105 (2014).

10. Sapranauskas, R. et al. The Streptococcus thermophilus CRISPR/Cas system provides immunity in Escherichia coli. Nucleic Acids Res. 39: 9275–9282 (2011).

11. Auer, T. O., Duroure, K., De Cian, A., Concordet, J. P. & Del Bene, F. Highly efficient CRISPR/Cas9-mediated knock-in in zebrafish by homology-independent DNA repair. Genome Res. 24: 142–153 (2014).

12. Hisano, Y. et al. Comprehensive analysis of sphingosine-1-phosphate receptor mutants during zebrafish embryogenesis. Genes Cells 20: 647–658 (2015).

13. Kimura, Y., Hisano, Y., Kawahara, A. & Higashijima, S. Efficient generation of knock-in transgenic zebrafish carrying reporter/driver genes by CRISPR/Cas9-mediated genome engineering. Sci. Rep. 4: 6545 (2014).

14. Li, R. et al. Antinociceptive effects of dexmedetomidine via spinal substance P and CGRP. Transl. Neurosci. 6: 259–264 (2015).

15. Shah, A. N., Davey, C. F., Whitebirch, A. C., Miller, A. C. & Moens, C. B. Rapid reverse genetic screening using CRISPR in zebrafish. Nat. Methods 12: 535–540 (2015).

16. Saxena, S. et al. Proteomic analysis of zebrafish caudal fin regeneration. Mol. Cell Proteomics 11: M111.014118 (2012).

17. Hwang, W. Y. et al. Efficient genome editing in zebrafish using a CRISPR-Cas system. Nat. Biotechnol. 31: 227–229 (2013).

18. Purushothaman, S. et al. Transcriptomic and proteomic analyses of Amphiura filiformis arm tissue undergoing regeneration. J. Proteomics 112: 113–124 (2015).

